# Chromatin adaptors and TOPBP1 condensates cooperate to organize ATM signaling

**DOI:** 10.64898/2026.08.21.746129

**Authors:** Clémence Mooser, Jihane Basbous, Nathalie Varisco, Tom Egger, Mònica Torres Esteban, Johannes Leyrer, Andrea Hänel, Vira Chea, Alain Jeanrenaud, Angelos Constantinou, Manuel Stucki

## Abstract

DNA damage response proteins frequently accumulate in biomolecular condensates, yet how these structures cooperate with classical adaptor-mediated recruitment mechanisms to organize DNA damage signaling remains poorly understood. Here, we identify Treacle and the ATM adaptor NBS1 as prominent components of TOPBP1 condensates and show that these structures activate ATM signaling in the absence of DNA damage. At rDNA breaks, Treacle recruits NBS1 and TOPBP1 through genetically separable interaction modules. Acute protein degradation further revealed that Treacle is required for condensate assembly, whereas TOPBP1 remains continuously required for condensate maintenance. Finally, we show that efficient ATM accumulation at IR-induced DNA double-strand breaks similarly depends on both NBS1 and TOPBP1, indicating that this two-component mechanism is not restricted to nucleolar DNA damage. Together, our findings support a two-component model in which adaptor proteins provide molecular specificity, whereas TOPBP1 condensates create the spatial organization required for robust ATM signaling.

## Introduction

The DNA damage response (DDR) in mammalian cells is coordinated by three phosphoinositide 3-kinase-related kinases (PIKKs): DNA-PKcs, ATM and ATR. While DNA-PK is predominantly implicated in the regulation of DNA double-strand break (DSB) repair by the non-homologous end joining pathway, ATM and ATR have a much broader spectrum of cellular functions and are also involved in cell signaling events that not only regulate DNA repair, but also cell cycle checkpoint activation, transcription, apoptosis and senescence (reviewed in Blackford and Jackson, 2017). ATM is thought to mostly respond to DSBs, while ATR is activated by a broader spectrum of DNA lesions. Notably, all three PIKKs require specific adaptors for their recruitment and activation at sites of DNA lesions (Falck et al., 2005).

How PIKK signaling is spatially organized at sites of DNA damage is not yet understood in molecular detail but biomolecular condensates have emerged as a potential shared organizing principle underlying activation of PIKK-family members. Rather than relying solely on diffusion-limited encounters between kinases and their activators and substrates, phase-separated assemblies concentrate signaling components at sites of damage, converting a stochastic search into a compartmentalized reaction. This principle is best established for the ATR kinase, for which its principal activator, TOPBP1, was shown to form biomolecular condensates via its intrinsically disordered ATR-activation domain, thus facilitating phosphorylation and activation of its downstream target CHK1 (Frattini et al., 2021; Egger et al., 2024). ATM activation has also been reported to be condensate-dependent, though through a distinct scaffold (Wang et al., 2022). These findings establish condensation as a potentially general mechanism for threshold-dependent activation of PIKKs, with TOPBP1 condensation specifically defined, to date, as an ATR-selective switch. Whether or not TOPBP1 can influence the activation of other PIKKs has not yet been investigated.

TOPBP1 is a versatile multivalent adaptor protein with various functions in DNA replication, DNA damage signaling and chromosome stability maintenance. Its many protein-protein interaction domains and motifs make TOPBP1 an ideal candidate for a coordinator and spatio-temporal organizer of multi-factor complexes. Thus, TOPBP1 repeatedly appears wherever large multi-protein signaling assemblies have to be built, for example during DNA replication initiation (Lim et al., 2023), ATR activation (Wardlaw et al., 2014), chromosome stability maintenance during mitosis (Adam et al., 2021; De Marco Zompit et al., 2022) or rDNA repair in the nucleoli (Velichko et al., 2019; Mooser et al., 2020). The latter is particularly interesting in the context of signaling competent biomolecular condensates because TOPBP1 interacts with the nucleolar phosphoprotein TCOF1/Treacle, thus regulating transcriptional repression in response to rDNA damage and nucleolar segregation, a process whereby rDNA repeats and associated proteins migrate from within the nucleoli in nucleolar caps, focal regions in the nucleolar periphery (Mooser et al., 2020; Petrova et al., 2026). Previous work showed that enforced TOPBP1 assembly, either through overexpression or induced multimerization, is sufficient to trigger biomolecular condensate formation and ATR activation in the absence of DNA damage (Sokka et al., 2015; Mooser et al., 2020; Frattini et al., 2021; Egger et al., 2024; Petrova et al., 2026). These observations indicate that signaling-competent TOPBP1 assemblies can form independently of DNA lesions. Whether or not Treacle-TOPBP1 biocondensates can also organize ATM signaling has not been explored. But since TOPBP1 interacts with proteins involved in both ATR and ATM signaling pathways, it is an attractive candidate to coordinate PIKK signaling more broadly.

Here, we investigate whether TOPBP1 condensates contribute to ATM signaling. We show that TOPBP1 condensates contain Treacle and the ATM adaptor NBS1 and promote ATM-dependent signaling. Furthermore, our findings uncover distinct roles for Treacle and TOPBP1 in condensate assembly and maintenance and suggest that TOPBP1-dependent signaling compartments may also function at canonical DSBs.

## Results

### TOPBP1 drives the formation of nuclear condensates containing Treacle, NBS1 and ATM

To define the molecular composition of TOPBP1 condensates, we combined optogenetic induction of TOPBP1 assembly with TurboID proximity labeling and quantitative mass spectrometry (Figure 1A). While many known TOPBP1 interactors were detected irrespective of light exposure, a small subset of proteins was preferentially enriched following condensate induction (Supplementary Table S1). Among the proteins preferentially labeled in the proximity of TOPBP1 were the nucleolar adaptor Treacle, the DNA repair factor BRCA1, and, unexpectedly, the ATM adaptor NBS1. Western blotting of Streptavidin-precipitated proteins confirmed the preferential proximity labeling of Treacle and NBS1 in cells exposed to blue light (Figure 1B). Interestingly, while proximity labeled ATM was not enriched in cells exposed to blue light, the active form of ATM, auto-phosphorylated on Ser1981 (ATM pS1981) was, suggesting that opto-TOPBP1 condensate formation may induce ATM autophosphorylation in the absence of DNA damage (Figure 1B).

**Figure 1.**
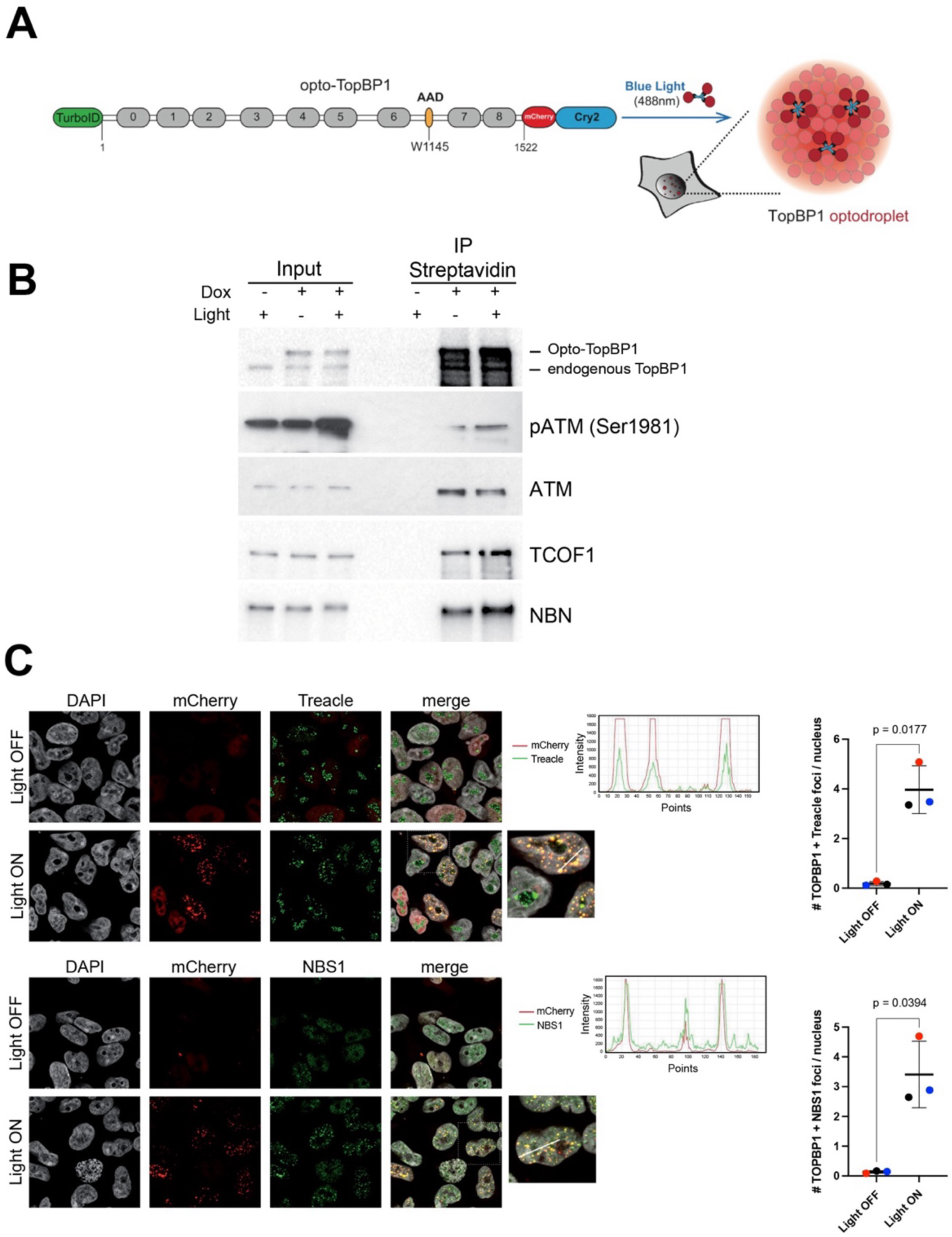
TOPBP1 drives the formation of nuclear condensates containing Treacle, NBS1 and ATM. (A) Schematic representation of the TopBP1 optogenetic platform. (B) Streptavidin pull-downs of proteins biotinylated by TurboID-TOPBP1 were probed for the indicated proteins using immunoblotting. When indicated (+), expression of TurboID-TOPBP1 was induced with 2 µg/mL doxycycline. When indicated (+), cells were exposed to blue light. (C) Immunofluorescence staining with the indicated antibodies of opto-TOPBP1 expressing cells activated by light. Line scans show co-localization. Graphs show the mean number of TOPBP1/Treacle and TOPBP1/NBS1 co-localizing foci per cell, respectively, of three independent experiments (color coded). Lines represent the mean of means and error bars the standard deviation. Statistical significance was calculated with paired two-tailed t-tests. Actual p-values are indicated (n=3, α = 0.05).

Immunofluorescence microscopy confirmed that light-induced TOPBP1 condensates efficiently recruited both Treacle and NBS1 (Figure 1C). Consistent with previous studies, depletion of Treacle largely prevented condensate formation (Figure S1A,B) (Mooser et al., 2020; Petrova et al., 2026).

Together, these findings identify Treacle and NBS1 as components of TOPBP1 condensates and suggest that these structures may support ATM signaling in addition to their established role in ATR activation.

### TOPBP1 condensates activate ATM signaling in the absence of DNA damage

Because optogenetic TOPBP1 condensation activates ATR signaling in the absence of DNA damage (Frattini et al., 2021), we asked whether it could similarly activate the ATM pathway. Light-induced TOPBP1 condensation rapidly triggered ATM autophosphorylation on Ser1981 followed by phosphorylation of CHK2, KAP1 and p53, demonstrating activation of canonical ATM signaling (Figure 2A,B; Figure S2). ATM inhibition abolished CHK2 and KAP1 phosphorylation without affecting CHK1, confirming that these signaling events were ATM-dependent (Figure 2B). ATR inhibition also suppressed CHK2 and KAP1 phosphorylation, consistent with the previously described requirement of ATR activity for optogenetic TOPBP1 condensation (Frattini et al., 2021).

**Figure 2.**
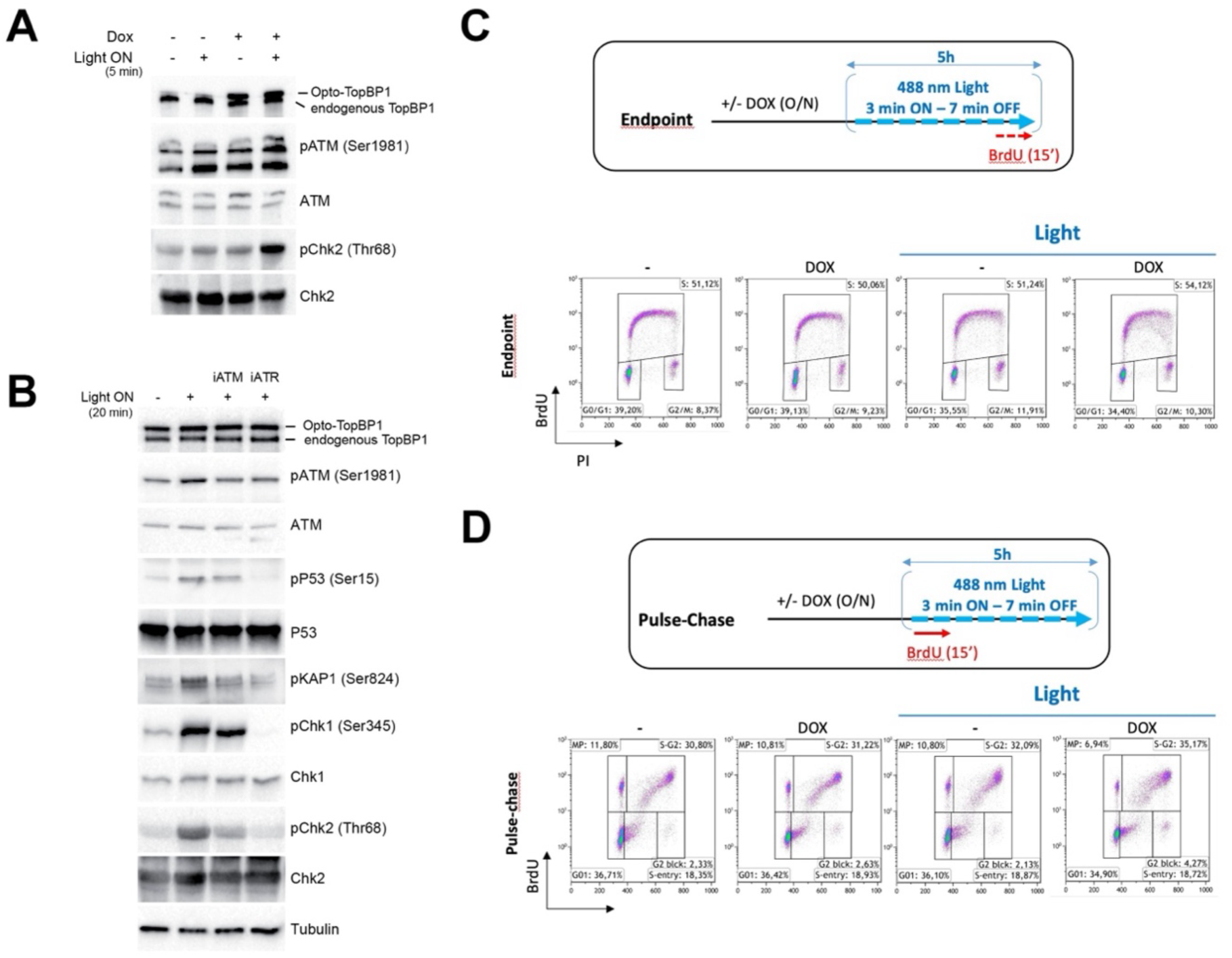
TOPBP1 condensates activate ATM signaling in the absence of DNA damage. (A) The indicated proteins were probed using immunoblotting after DOX induction of the expression of opto-TOPBP1 and light activation, as indicated. (B) The indicated proteins were probed using immunoblotting of cells expressing opto-TOPBP1 after light activation, as indicated. Cells were pre-incubated with ATM inhibitors (iATM) or ATR inhibitors (iATR) as indicated. (C) Flow cytometry analysis of cells inducibly expressing opto-TOPBP1 with and without exposure to repeated cycles of blue light. When indicated (DOX) opto-TOPBP1 expression was induced. Cells were subjected to a 15 min BrdU pulse at the end of the light exposure. Percentage of cells in the cell cycle phases G0/G1, S and G2/M are indicated. Top panel shows a schematic timeline of the experiment. (D) Flow cytometry analysis of cells inducibly expressing opto-TOPBP1 with and without exposure to repeated cycles of blue light. When indicated (DOX) opto-TOPBP1 expression was induced. Cells were subjected to a 15 min BrdU pulse before light exposure (pulse chase). Percentage of cells in the cell cycle phases G0/G1, S and G2/M are indicated. MP stands for ‘mitosis passed’ and represents cells that successfully completed mitosis and therefore present with 2n DNA, labeled with BrdU. Top panel shows a schematic timeline of the experiment.

To determine whether sustained TOPBP1 condensation affects cell-cycle progression, we repeatedly induced condensate formation over several hours before monitoring DNA synthesis by BrdU incorporation. Flow cytometry analysis revealed a subtle collapse of BrdU signals in cells expressing opto-TOPBP1 that were exposed to the light. No such effect was observed in cells that did not express opto-TOPBP1 or were not exposed to blue light (Figure 2C). Pulse-chase analysis revealed accumulation in G2 and a reduction in the fraction of cells that had progressed through mitosis (Figure 2D; MP fraction, for “mitosis passed”).

Together, these findings demonstrate that TOPBP1 condensates are functional signaling compartments that activate both ATM and ATR pathways and are sufficient to elicit checkpoint responses in the absence of exogenous DNA damage.

### Treacle coordinates ATM recruitment through genetically separable NBS1 and TOPBP1 interaction modules

To determine whether TOPBP1 also contributes to ATM recruitment at physiological DNA lesions, we induced targeted DSBs within the rDNA repeats by I-PpoI mRNA transfection and monitored protein accumulation at nucleolar caps by immunofluorescence microscopy. Specificity of the ATM and ATR antibodies was confirmed by siRNA-mediated depletion of the respective proteins (Figure S3A,B). As expected, loss of NBS1 abolished ATM, but not ATR, recruitment to nucleolar caps following rDNA break induction (Figure 3A). Reconstitution of U2OS ΔNBS1 cells with wild type NBS1, but not with a C-terminal deletion mutant lacking the ATM interaction motif, rescued ATM recruitment (Figure 3B). These findings confirm NBS1 as an ATM-specific adaptor. Surprisingly though – but consistent with the results obtained on the optogenetic platform – TOPBP1 depletion not only abolished the recruitment of ATR to sites of rDNA breaks, but it also abrogated the recruitment of ATM to nucleolar caps (Figure 3C). These findings identify TOPBP1 as an unexpected regulator of ATM recruitment at endogenous DNA breaks, consistent with our observations in the optogenetic system.

**Figure 3.**
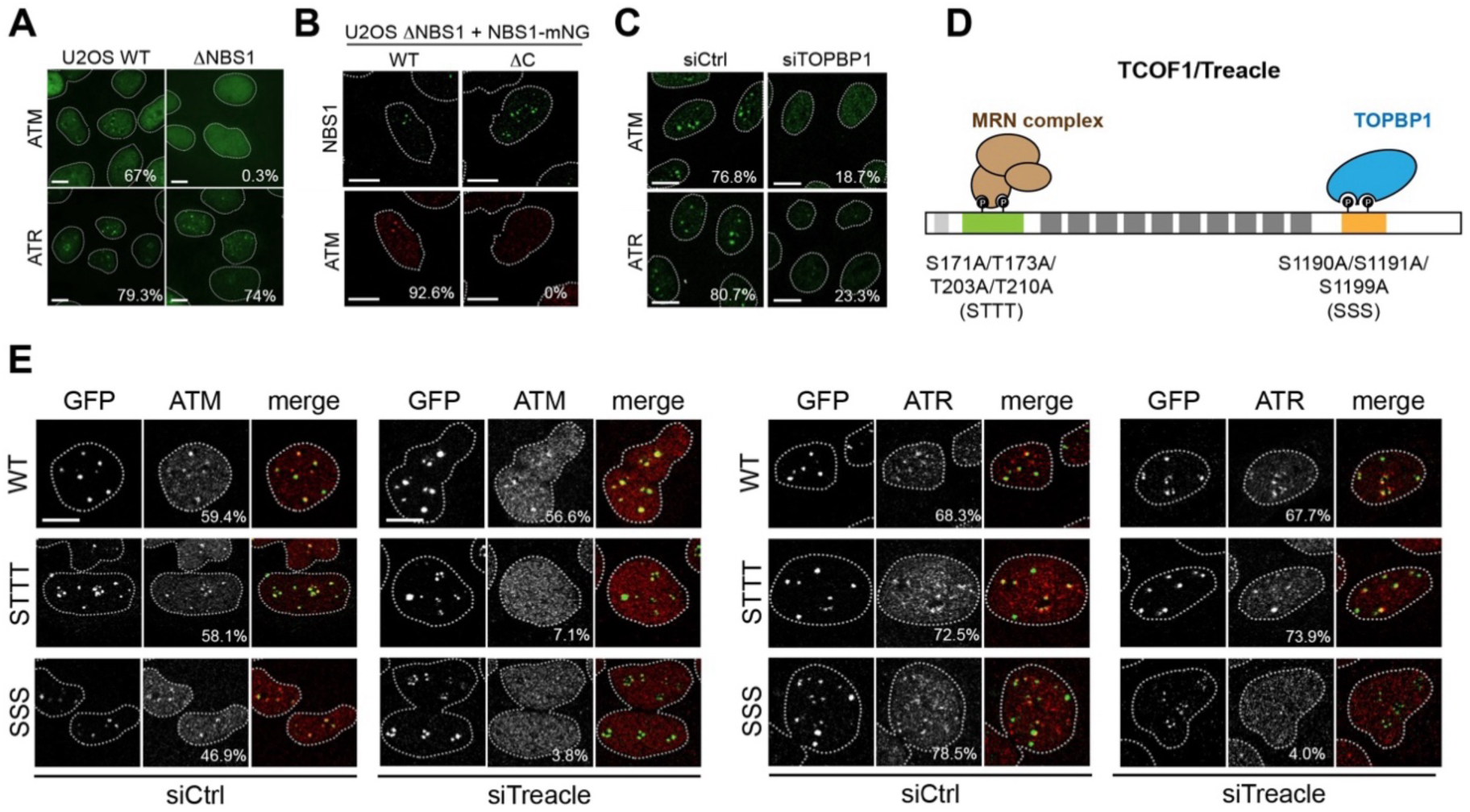
Treacle coordinates ATM recruitment through genetically separable NBS1 and TOPBP1 interaction modules. (A) ATM and ATR localization in U2OS ΔNBS1 cells and control cells 2h after I-PpoI transfection. Percentage of cells with > 2 ATM or ATR caps are indicated. (B) ATM localization in U2OS ΔNBS1 cells stably transfected with full-length mNeonGreen (mNG) tagged NBS1 wild type (WT) and C-terminal deletion mutant (ΔC). Percentage of cells with > 2 ATM caps are indicated. (C) ATM and ATR localization in TOPBP1 depleted U2OS cells and control cells 2h after I-PpoI transfection. Percentage of cells with > 2 ATM or ATR caps are indicated. (D) Schematic showing the layout of NBS1 and TOPBP1 binding regions in Treacle. The phosphorylation sites required for NBS1 and TOPBP1 binding are indicated. (E) ATM and ATR localization 2 h after I-PpoI transfection in GFP-Treacle-expressing cell lines (WT or STTT and SSS mutants, respectively) after treatment with control siRNA (left) and siTreacle (right). Percentage of cells with > 2 ATM or ATR caps are indicated. Panels A-C and E: one representative of two independent experiments is shown. Scale bar = 10 µm.

To distinguish the contributions of the NBS1- and TOPBP1-binding sites within Treacle, we generated U2OS cells stably expressing siRNA-resistant wild-type Treacle or mutants selectively disrupting either interaction (Larsen et al., 2014; Mooser et al., 2020) and assessed in each cell line how well ectopic expression of Treacle could rescue the effect of endogenous Treacle downregulation (Figure 3D). As expected, depletion of endogenous Treacle by siRNA transfection robustly abrogated both ATM and ATR accumulation in nucleolar caps (Figure S3C). Wild-type Treacle efficiently rescued ATM recruitment following depletion of endogenous Treacle, whereas the NBS1-binding mutant (STTT) failed to do so (Figure 3E). In contrast, ATR recruitment remained unaffected in STTT cells. Disruption of the TOPBP1-binding site (SSS), however, abolished recruitment of both ATM and ATR (Figure 3E). These data thus show that Treacle contains two genetically separable recruitment modules, one for NBS1, one for TOPBP1. Both are required for ATM recruitment, but only one is required for ATR recruitment. ATM and ATR recruitment occurred independently of γH2AX-MDC1 signaling (Figure S3D), indicating that nucleolar recruitment relies on a distinct upstream mechanism.

### Recruitment of the Treacle-NBS1-TOPBP1 complex is required for productive ATM signaling

To determine whether the Treacle-TOPBP1 axis is required for ATM signaling following rDNA break induction, we monitored phosphorylation of the downstream ATM and ATR targets CHK2 and CHK1, respectively. Depletion of TOPBP1 by siRNA led to significant reduction of both CHK1 and CHK2 phosphorylation (Figure 4A). Loss of Treacle expression had an intermediate effect, with both CHK1 and CHK2 phosphorylation being reduced, but not fully abrogated (Figure 4B). These findings indicate that the defects in ATM recruitment are accompanied by impaired downstream signaling.

**Figure 4.**
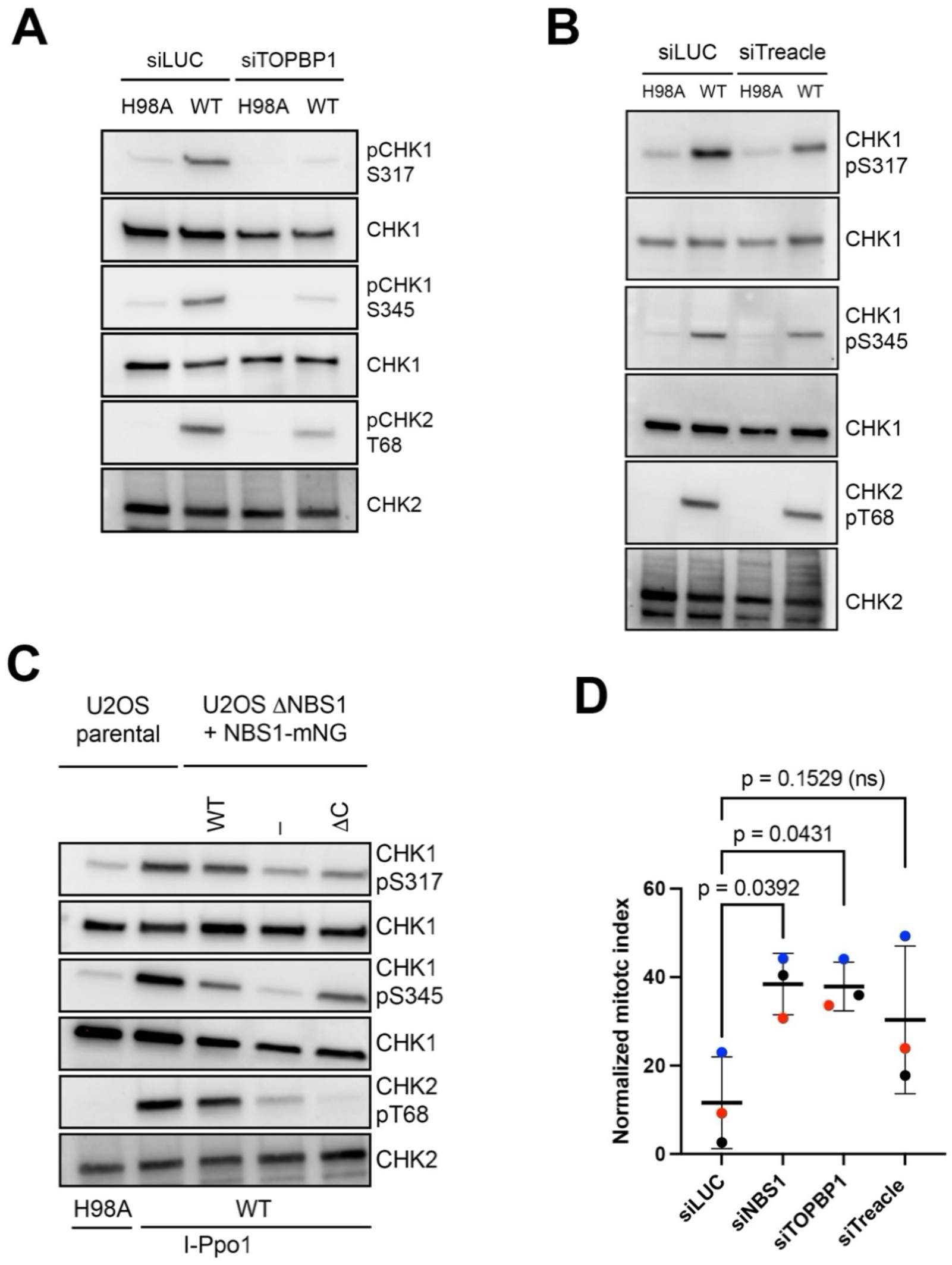
Recruitment of the Treacle-NBS1-TOPBP1 complex is required for productive ATM signaling. (A) Western blots of total cell extracts of U2OS cells, control depleted (siLUC) or depleted of TOPBP1 (siTOPBP1), and transfected with I-PpoI wild type (WT) and catalytic mutant H98A, respectively. Blots were stained with the indicated antibodies. (B) Western blots of total cell extracts of U2OS cells, control depleted (siLUC) or depleted of Treacle (siTreacle), and transfected with I-PpoI wild type (WT) and catalytic mutant H98A, respectively. Blots were stained with the indicated antibodies. (C) Western blots of total cell extracts of U2OS ΔNBS1 cells, either stably complemented with empty vector (–), full-length mNeonGreen tagged NBS1 wild type (WT) and mNeonGreen tagged NBS1 C-terminal ATM interaction mutant (ΔC), and transfected with I-PpoI wild type (WT) and catalytic mutant H98A, respectively. Blots were stained with the indicated antibodies. (D) Percentage of mitotic cells in response to I-PpoI transfection in control depleted cells (siCtrl), NBS1 (siNBS1), TOPBP1 (siTOPBP1) and Treacle (siTreacle) depleted cells, respectively, normalized in each case to untreated cells without I-PpoI transfection. Dots represent three independent experiments (color coded), the black line indicates the mean and the error bars the standard deviation. Statistical significance was tested by ordinary one-way ANOVA with Dunnett’s multiple comparison test. Actual p values are indicated (n = 3; α = 0.05; ns = non-significant).

In U2OS ΔNBS1 cells, both CHK1 phosphorylation on Ser117 and Ser345 and CHK2 phosphorylation on Thr68 were severely affected (Figure 4C). Complementation with wild type NBS1 mostly rescued the phosphorylation defect. In contrast, expression of the ΔC mutant restored CHK1 phosphorylation but failed to rescue CHK2 phosphorylation, demonstrating that the direct interaction between NBS1 and ATM is specifically required for ATM signaling.

To determine whether these signaling defects compromise checkpoint function, we measured the mitotic index following rDNA break induction. Control cells exhibited a marked reduction in mitotic cells, consistent with activation of a robust G2/M checkpoint. In contrast, depletion of NBS1 and TOPBP1 significantly attenuated this response, indicating defective checkpoint activation (Figure 4D). Depletion of Treacle also attenuated the response, but the difference was not statistically significant.

Together, these findings demonstrate that recruitment of the Treacle-NBS1-TOPBP1 complex to rDNA breaks is required not only for ATM accumulation, but also for productive ATM-dependent checkpoint signaling.

### TOPBP1 maintains nucleolar condensates after Treacle-dependent assembly

To distinguish the roles of Treacle and TOPBP1 in condensate maintenance, we generated DLD1 cells expressing endogenous dTag-tagged TCOF1 alleles and used an analogous published dTag-tagged TOPBP1 cell line (Nabet et al., 2018; Trivedi et al., 2023) (Figure S4A,B). Acute addition of dTAG-13 efficiently degraded Treacle and TOPBP1 with half-lives of approximately 26 min and 14 min, respectively (Trivedi et al., 2023) (Figure S4A-C).

Interestingly, downregulation of Treacle after I-PpoI-induced nucleolar segregation did not affect TOPBP1 and NBS1 localization within nucleolar caps. In addition, nucleolar segregation was maintained for the duration of the experiment (Figure 5A,B). In contrast, downregulation of TOPBP1 at the same timepoint led to rapid collapse of Treacle- and NBS1-positive nucleolar caps and a reversion of nucleolar segregation and migration of Treacle inside of the nucleoli (Figure 5C). This progressive loss of NBS1-positive caps reached statistical significance after 2 and 3 h of dTAG-13 treatment (Figure 5D).

**Figure 5.**
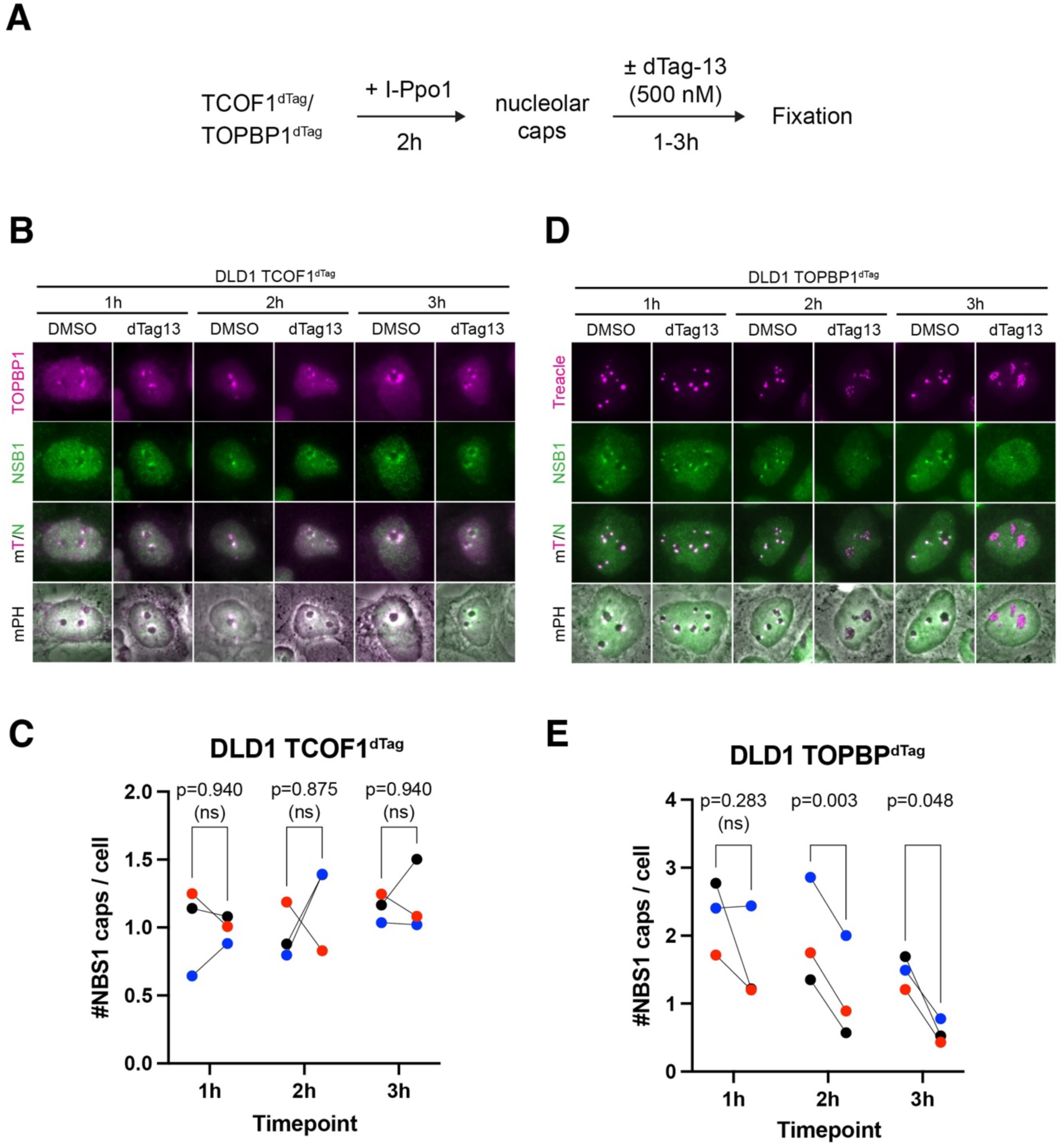
TOPBP1 maintains nucleolar condensates after Treacle-dependent assembly. (A) Experimental scheme. rDNA double-strand breaks and nucleolar caps formation were induced with *in vitro*-transcribed I-PpoI mRNA for 2 h, followed by treatment with dTag-13 or DMSO for an additional 1, 2, or 3 h before fixation. (B) Representative images TCOF1^dTag^ cells treated and stained as indicated. (C) Quantification of the data in (B). Dots represent the mean number of NBS1-positive nucleolar caps per cell from three independent experiments (color-coded). DMSO- and dTAG-13-treated samples from the same experiment are connected by lines. Statistical significance was assessed using paired two-tailed t-tests with Holm– Šídák correction for multiple comparisons across the three time points. Adjusted p-values are indicated. (α = 0.05; ns = non-significant). (D) Representative images TOPBP1^dTag^ cells treated and stained as indicated. (E) Quantification of the data in (D). Dots represent the mean number of NBS1-positive nucleolar caps per cell from three independent experiments (color-coded). DMSO- and dTAG-13-treated samples from the same experiment are connected by lines. Statistical significance was assessed using paired two-tailed t-tests with Holm– Šídák correction for multiple comparisons across the three time points. Adjusted p-values are indicated. (a = 0.05; ns = non-significant).

Together, these findings demonstrate that Treacle and TOPBP1 fulfill distinct functions during nucleolar condensate formation, with Treacle acting during condensate establishment and TOPBP1 remaining continuously required to maintain the assembled signaling compartment.

### The two-component mechanism of ATM accumulation extends to canonical DSBs

To determine whether the role of TOPBP1 in ATM recruitment extends beyond nucleolar DNA damage, we induced randomly distributed DSBs by ionizing radiation (IR). The validated ATM antibody detected prominent ATM irradiation-induced foci (IRIF) that colocalized with γH2AX and, consistent with TOPBP1, were most prominent in G1 cells (Cescutti et al., 2010; Leimbacher et al., 2019) (Figure S5A,B).

Depletion of both NBS1 and TOPBP1 by siRNA abrogated ATM IRIF, thus reflecting the same ATM accumulation dependency pattern observed in response to targeted rDNA break induction (Figure 6B and S5C). In addition, depletion of the upstream regulators of TOPBP1 recruitment, MDC1 and 53BP1, by siRNA abrogated ATM IRIF (Figure 6B and S5C). Thus, ATM accumulation outside the nucleolus also requires both its canonical NBS1 adaptor and TOPBP1.

**Figure 6.**
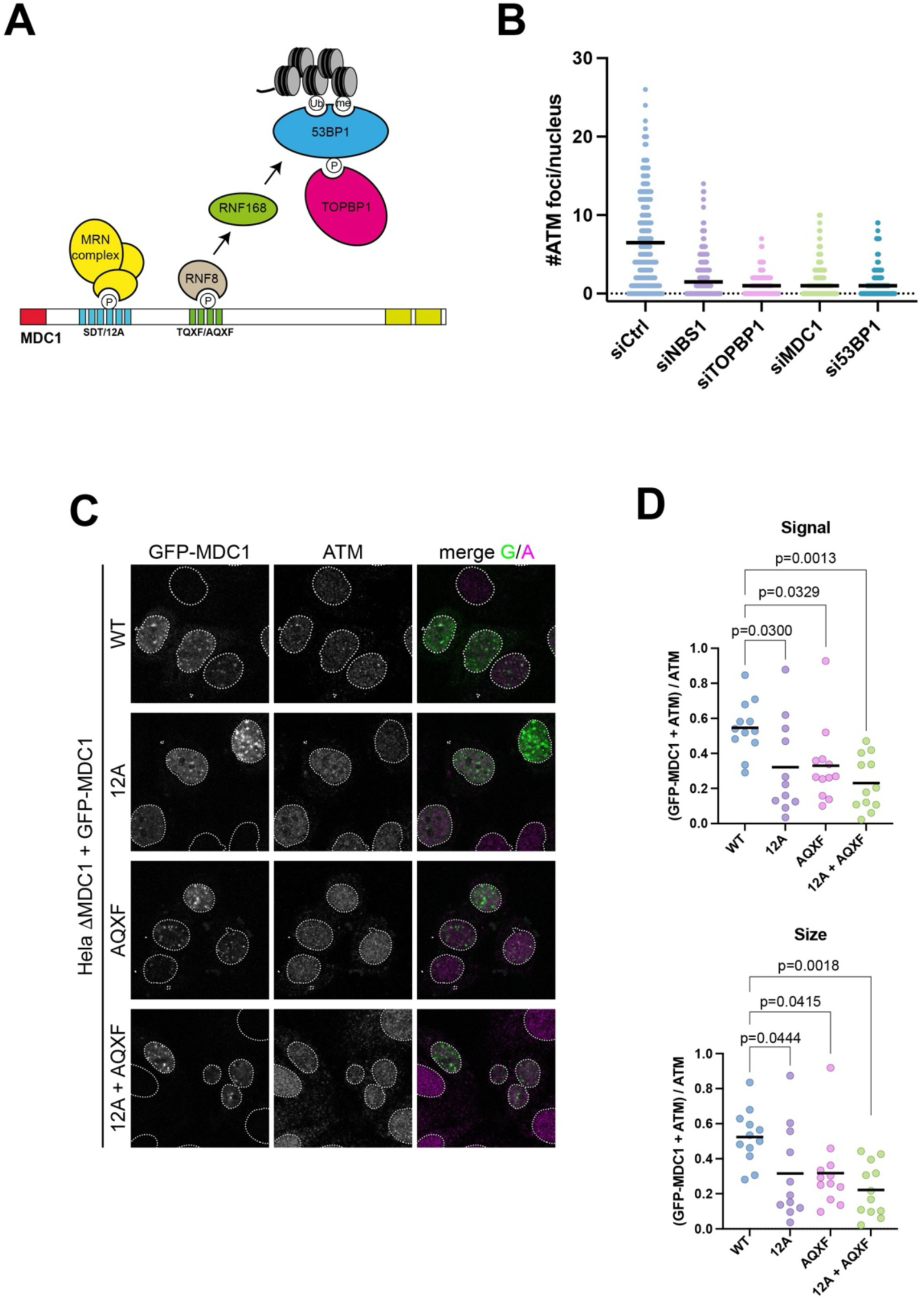
The two-component mechanism of ATM accumulation extends to canonical DSBs. (A) Schematic representation of the two phoshorylation site clusters in MDC1 that mediate the accumulation of the MRN complex and TOPBP1 (via the RNF8-RNF168-53BP1 axis) at sites of IR-induced DSBs. (B) ATM foci formation in Hela cells either mock depleted (siCtrl) or depleted of NBS1 (siNBS1), TOPBP1 (siTOPBP1), MDC1 (siMDC1) and 53BP1 (si53BP1), respectively, 3h after exposure to 3 Gy of IR. Data from one of two independent experiments is shown. Each dot represents one cell; horizontal lines indicate the mean. (C) GFP-MDC1 and ATM localization in HeLa ΔMDC1 cells expressing GFP-MDC1 wild type (WT), SDT repeat mutant (12A), TQXF cluster mutant (AQXF) and the combination mutant (12A + AQXF). (D) Quantitative analysis of GFP-MDC1 and ATM colocalization by SQUASSH. Upper graph: object signal colocalization (sum of all pixel intensities in one channel where objects overlap with objects from the other channel divided by total sum of object pixel intensities). Lower graph: object size colocalization (area of object overlap divided by total object area). Each data point represents one cell (n = 12); bars represent the mean. Statistical significance was tested with ordinary one-way ANOVA and Dunnett’s multiple comparison. Actual p values are indicated (α = 0.05).

Outside the nucleolus, recruitment of NBS1 and TOPBP1 is coordinated by two distinct phosphorylation-dependent interaction modules within MDC1: the SDT repeats recruit NBS1, whereas the TQXF cluster promotes TOPBP1 accumulation through the RNF8-RNF168-53BP1 route (Spycher et al., 2008; Melander et al., 2008; Wu et al., 2008; Chapman and Jackson, 2008; Mailand et al., 2007; Kolas et al., 2007; Huen et al., 2007; Doil et al., 2009; Stewart et al., 2009; Cescutti et al., 2010; Leimbacher et al., 2019) (Figure 6A).

To determine whether both recruitment modules contribute to ATM accumulation we knocked-out endogenous MDC1 in Hela cells using CRISPR-Cas9 (henceforth termed ΔMDC1), followed by transient transfection of the knock-out cells with GFP-tagged full-length MDC1 constructs. WT and phosphorylation-deficient MDC1 mutants accumulated efficiently at IRIF. However, ATM recruitment was restored only by WT MDC1, whereas disruption of either the SDT repeats or the TQXF cluster significantly impaired ATM accumulation (Figure 6C,D).

Together, these findings indicate that efficient ATM accumulation at both nucleolar and canonical DNA double-strand breaks requires two complementary recruitment mechanisms: direct binding of ATM to NBS1 and TOPBP1-dependent assembly of higher-order signaling compartments.

## Discussion

This study identifies TOPBP1 as a previously unrecognized organizer of ATM signaling. While TOPBP1 has long been appreciated as the principal activator of ATR, our findings indicate that it also contributes to efficient ATM accumulation and signaling at rDNA breaks, thereby extending the role of TOPBP1 from ATR activation to the broader organization of ATM and ATR signaling. Our data indicate that the same organizational principle likely extends beyond the nucleolus, since efficient ATM accumulation at IR-induced DSBs also requires TOPBP1. However, unlike at rDNA breaks, our current evidence outside the nucleolus is limited to ATM recruitment rather than downstream signaling.

Efficient ATM assembly requires two fundamentally different organizational principles. The first one is mediated by the canonical ATM adaptor NBS1. It provides molecular specificity through direct ATM binding via its C-terminal ATM interaction motif (Falck et al., 2005; Warren and Pavletich, 2022). TOPBP1, for which no convincing direct interaction with ATM has yet been reported, instead appears to provide spatial organization through higher-order assembly. This raises an obvious question: if NBS1 already recruits ATM to sites of DNA damage, why is TOPBP1 required?

The conventional view of PIKK activation involves their binding to specific aberrant DNA structures commonly understood as the upstream signals of the DDR. But a growing body of work has challenged the view that DNA damage signaling depends exclusively on aberrant DNA structures. Together, these studies established that artificial chromatin tethering, overexpression or induced oligomerization of TOPBP1 is sufficient to trigger ATR signaling independently of DNA damage (Lindsey-Boltz and Sancar, 2011; Sokka et al., 2015; Frattini et al., 2021; Petrova et al., 2026). Interestingly, the latter two responses are dependent on Treacle, and, more precisely, on the TOPBP1 interaction sites in Treacle, indicating that they also require a chromatin tethering function (Figure S1D) (Mooser et al., 2020; Petrova et al., 2026). Such DNA damage independent TOPBP1 conglomerates display characteristics of liquid phase separated biomolecular condensates and are capable of inducing ATR signaling in the absence of DNA damage (Frattini et al., 2021; Petrova et al., 2026). In this work, we show that these TOPBP1 condensates can also induce an ATM signaling cascade in the absence of DSBs, indicating that they are also implicated in the organization of ATM signaling.

The distinction between condensate assembly and condensate maintenance has received relatively little attention in the field. Most studies implicitly treat condensate assembly and maintenance as the same process. Here we present evidence that these two processes are genetically separable. Treacle mainly acts as a nucleation factor and once the condensates have formed, it becomes dispensable. On the other hand, without TOPBP1, the condensates rapidly collapse. Our findings therefore identify Treacle and TOPBP1 as functionally distinct components of the signaling compartment: Treacle initiates DNA damage-induced condensate assembly, whereas TOPBP1 maintains the assembled compartment.

In summary, our work suggests the following refined model for activation and maintenance of ATM signaling: First, chromatin-bound adaptor proteins recruit the MRN complex and ATM to sites of DNA damage through NBS1. These adaptors simultaneously nucleate TOPBP1 condensate assembly, which provides the higher-order organization required for efficient ATM accumulation and formation of a stable signaling compartment.

Classical adaptor proteins and biomolecular condensates should not be viewed as alternative mechanisms for assembling DNA damage signaling complexes. Instead, our findings suggest that they fulfill complementary functions. Chromatin-bound adaptors provide molecular specificity by recruiting the appropriate signaling factors to sites of DNA damage, whereas TOPBP1-dependent condensates organize these factors into higher-order signaling compartments that enable efficient and sustained DNA damage signaling. Importantly, condensate formation itself remains dependent on chromatin-bound adaptor proteins, indicating that higher-order organization builds upon, rather than replaces, classical adaptor-mediated recruitment. Although our study focuses on ATM signaling, the complementary roles of adaptors and condensate-forming scaffold proteins may represent a more general organizational principle for the assembly of dynamic signaling compartments.

## Acknowledgements

We thank Don Cleveland, Steve Jackson and Brian McStay for providing valuable reagents. Imaging was performed with equipment maintained by the Centre for Microscopy and Image Analysis, University of Zürich. Cell sorting was carried out by the Flow Cytometry Core Facilities at the University of Zürich. Mass spectrometry experiments were carried out using the facilities of the Montpellier Proteomics Platform (PPM, BioCampus Montpellier). The Basbous lab is supported by the French Agence Nationale de la Recherche ANR BioTop (AAPG2021) and the Fondation ARC (projet ARC 2024 PJA3). The Stucki lab is supported by two project grants from the Swiss National Foundation (31003A_163141 and 310030_189141) and by the Kanton of Zürich. We dedicate this work to the memory of Angelos Constantinou, whose scientific insight and mentorship were instrumental in initiating this project.

## Author Contributions

The project was conceived and supervised by A.C., J.B. and M.S. Biochemical and cell biological experiments were planned and carried out and data were analyzed by C.M., J.B., N.V., T.E., M.T.E., J.L., A.H. and V.C. Reagents were generated by A.J. The paper was written by C.M., J.B. and M.S. with contributions from the other authors.

## Methods

### Cell culture and drug treatments

U2OS (human osteosarcoma cell line), HeLa (human cervical carcinoma cell line, donor: Henrietta Lacks), RPE-1 hTERT (immortalized human retinal pigmented cell lines) and 293FT (transformed human embryonal kidney cell line) were grown in a sterile culture environment in Dulbecco’s modified Eagle medium (DMEM Gibco^TM^, 61965026), supplemented with 10% fetal calf serum (PAN Biotech, P40-37500), 2mM L-glutamine, and penicillin-streptomycin antibiotics. DLD-1 cells were cultured in RPMI-1640 medium (PAN-Biotech, P04-18047) supplemented with 10% fetal calf serum (PAN-Biotech, P40-37500) and 1% penicillin–streptomycin (Gibco™, 15240-062). All cell lines were maintained under standard cell culture conditions in a CO_2_ incubator (37°C, 5% CO_2_). U2OS cell lines stably transfected with NBS1-mNG expression constructs were cultured in the presence of 1 µg/ml Puromycin (Sigma). U2OS cell lines stably transfected with MDC1-GFP and TOPBP1-GFP expression constructs were cultured in the presence of 400 µg/ml Geneticin (G418 sulfate, Sigma). U2OS cell lines stably transfected with Treacle-GFP expression constructs were cultured in the presence of 200 µg/ml Zeocin and 10 µg/ml Blasticidin S (Invitrogen). The following compounds were used at the indicated final concentrations: ATM inhibitor KU-55933 (5 µM) (Selleckchem), ATR inhibitor VE-821 (5 µM) (Selleckchem), dTag-13 (500 nM). All cell lines were regularly tested for mycoplasma contamination. For generation of random DSBs, X-ray irradiation was performed with YXLON.SMART 160E-1.5 device (150kV, 6mA, YXLON International SA) delivering 11.8 mGy per second. Soft X-rays were largely filtered out with a 3-mm aluminium filter. 5 or 10 Gy were used for the experiments. Flp-In™ 293 T-REx cells were obtained from Thermo Fisher Scientific and cultured in DMEM-GlutaMAX (Merck-Sigma-Aldrich, #D5796) supplemented with 10% heat-inactivated fetal bovine serum (BioWest FBS, Eurobio Scientific, #S1810). Stable cell lines expressing TOPBP1 fused at its N-terminus to TurboID and at its C-terminus to the photoreceptor cryptochrome 2 (Cry2) and mCherry (opto-TOPBP1) were generated as previously described (Frattini et al., 2021). Opto-TOPBP1-expressing cell lines were maintained in DMEM supplemented with 10% FBS, 5 µg/mL blasticidin, and 50 µg/mL hygromycin. For experiments, cells were seeded in selection-free medium, with or without 2 µg/mL doxycycline (referred to as “DOX” in the figures) to induce expression of the opto-TOPBP1 construct.

### Cloning and mutagenesis

The pcDNA4-TO-strep-HA-GFP-Treacle construct for GFP-Treacle expression, the S171A/T173A/T203A/T219A (STTT) mutant derivative and the S1227A/S1128A/S1236A (SSS) mutant derivate have been described (Larsen et al., 2014; Mooser et al., 2020). The pcDNA5/FRT/TO-NBS1-mNG construction for mNEON-NBS1 expression and the R28A/K160M mutant derivative have been described (Mooser et al., 2020). Mutant derivate NBS1-MRE11 interaction mutant was generated by site directed mutagenesis using the QuickChange II XL Site-Directed Mutagenesis kit (Agilent Technologies, 200522). The sequences of the primers used for the mutagenesis are provided in the key resources table. The pcDNA5/FRT-TO-MDC1-HA-WT construct for GFP-MDC1 expression, the SDT and AQXF mutant derivate have been described (Leimbacher et al., 2019). Plasmid pIRES V5 I-PpoI and pIRES V5 I-PpoI (H98A) for I-PpoI mRNA purification (gift from Brian McStay) were described (van Sluis and McStay, 2015). The pCR2.1_DDTag_TCOF1_Puromycin and pCR2.1_DDTag_TCOF1_Blasticidin donor plasmids (Park et al., 2014) were converted into dTag constructs by introducing the P107L substitution (CCG to CTG) using the QuikChange II XL Site-Directed Mutagenesis Kit (Agilent, #200521) according to the manufacturer’s instructions. The sequences of the primers used for the mutagenesis are provided in the key resources table. For the MDC1 CRISPR/Cas9 targeting plasmid, a sense and an antisense gRNA targeting the FHA domain of MDC1 with minimal off-target impact were introduced in the All-In-One (AIO) CRISPR-Cas9^D10A^-mCherry plasmid (Addgene) described in (Chiang et al., 2016). The sequences of the primers used for the cloning of the sense and antisense gRNAs are provided in the key resources table.

### Opto-TOPBP1 activation

Flp-In 293 T-REx cell lines stably expressing recombinant opto-TOPBP1 proteins were seeded at approximately 70% confluency in DMEM supplemented with 10% FBS, without selection. Expression of opto-TOPBP1 was induced for 16 h by treatment with 2 µg/mL doxycycline. For light activation, cells were transferred to a custom-built illumination box equipped with an array of 24 LEDs emitting at 488 nm and delivering a light intensity of 10 mW/cm², as measured using a Thorlabs PM16-121 power meter. Cry2 oligomerization was induced using light–dark cycles with the timing indicated in the figures, consisting of 4 s of illumination followed by 10 s in the dark.

### Generation of cell lines

The U2OS ΔNBS1 cells expressing NBS1-mNG wild type and mutants were generated by co-transfection of pcDNA5/FRT/TO-NBS1-mNG with pBabePuro, followed by selection with 1 µg/ml Puromycin. The GFP-TOPBP1 cells expressing GFP-TOPBP1 wild type and mutants were generated by transfection of pIRESneo2-GFP-TOPBP1 WT or MUT, followed by selection with 400 µg/ml Geneticin. To isolate clonal cells lines with homogenous TOPBP1-GFP expressing and NBS1-mNG expression, cell pools were sorted using DB FACSAria III 4L cell sorter. To generate an MDC1 knock-out in HeLa cells, cells were transfected with the AIO CRISPR-Cas9^D10A^-mCherry_MDC1gRNA plasmid using jetOPTIMUS® DNA Transfection Reagent (Polyplus). After 48 hours, mCherry-positive single cells were sorted into 96-well plates using a DB FACSAria III 4L cell sorter. Wells containing single colonies were identified under a transluminescence microscope and progressively expanded to larger culture volumes. MDC1 knock-out was confirmed by western blotting of total cell extracts of single clones and by Sanger sequencing of a PCR-amplified genomic fragment. To generate endogenously tagged TCOF1^dTag^ cell line, DLD-1 cells (3 × 10^5^) were seeded in 6-well plates and co-transfected the following day with 1 µg each of pX330-TCOF1 (Park et al., 2014) and the puromycin- and blasticidin-resistant dTag donor plasmids. Cells were selected with 1 µg/mL puromycin and 10 µg/mL blasticidin, expanded for approximately 2 weeks, and single-cell clones were isolated by limiting dilution. Positive clones were subsequently maintained without selection.

### RNA transfection

The control (siCtrl) and siRNA against Treacle (siTreacle), TOPBP1 (siTOPBP1), MDC1 (siMDC1), 53BP1 (si53BP1), NBS1 (siNBS1) were obtained from Microsynth AG. The sequences of the siRNA are provided in the key resource table. siRNA transfection (10 picoM) was performed using Lipofectamine RNAiMAX (Invitrogen) in cells grown in twelve-well plate in DMEM supplemented with 10% FBS according to the manufacturer’s instruction. Cells were split on coverslips 24 hours after transfection and harvested 72 hours after transfection. I-PpoI mRNA generation and purification was performed as described (van Sluis and McStay, 2015). The vectors pIRES I-PpoI WT and H98A were linearized with NotI and transcribed in vitro using the MEGAscript T7 kit (Ambion) according to the manufacturer’s protocol. The I-PpoI mRNA was polyadenylated using the Poly(A) tailing kit (Ambion) according to the manufacturer’s protocol. I-PpoI mRNA transfection was done using the Lipofectamine MessengerMax Reagent (Invitrogen) according to the manufacturer’s protocol.

### SDS-Page and western blotting

SDS-page and western blotting were performed using 4-20% Mini-PROTEAN TGX Stain-free Precast Gels (Bio-Rad, 4568094) and Trans-Blot Turbo 0.2µm nitrocellulose membranes (Bio-Rad. 1704158). The following antibodies were used at the indicated dilutions: CHK1 pS317 (rabbit, Cell Signaling, 2344, 1/500), CHK1 pS345 (rabbit, Cell Signaling, 2348, 1/500), CHK1 (mouse, Santa Cruz Biotechnology, sc-8404, 1/1000), CHK2 pT68 (rabbit, Cell Signaling, 2661, 1/1000), CHK2 (rabbit, Cell Signaling, 2662, 1/1000), NBS1 (rabbit, Abcam, ab32794, 1/2000), TOPBP1 (rabbit, abcam, Ab2402, 1/1500), phospho-KAP1 (S824) (A300-767A, Bethyl, 1/1000), ATM pS1981 (rabbit, abcam, Ab81292, 1/1000), ATM (rabbit, Cell Signaling, 2873, 1/1000), P53 pS15 (rabbit, Cell signaling, 9284, 1/1000), α-Tubulin (mouse, Sigma-Aldrich, T5168, 1/10000), Treacle (Proteintech, 11003-1-AP; 1/1,000). SuperSignal^TM^ West Femto Maximum Sensitivity Substrate (Thermo Scientific^TM^, 34096) was used for blot developments and ChemiDoc MP Imaging System (Bio-Rad, 17001402) for image acquisition.

### Immunofluorescence

Cells were grown in DMEM on glass coverslips for 24 hours and fixed with either 4% buffered formaldehyde for 15 minutes at room temperature or ice-cold methanol for 10 minutes. Cells were permeabilized for 5 minutes in PBS containing 0.3% Triton x-100. Cells were blocked for 1 hour in blocking buffer (10% FBS in PBS or 5% BSA in PBS) followed by primary antibody incubation for 1 h at room temperature or overnight at 4°C. The coverslips were washed three times in PBS and the secondary antibody incubation was performed for 1 hour at room temperature in the dark (anti-rabbit, Invitrogen, A21235 1/1000 or anti-mouse, Invitrogen, A10042, 1/1000). Following the incubation, coverslips were washed and mounted on glass microscopy slides (Thermo Scientific, 630-1985, dimensions L76 X W26 mm) with VECTASHIELD® mounting medium containing 0.5 µg/ml 4’,6-diamidino-2-phenylindole dihydrochloride (DAPI) (Vector Laboratories, H-1200). The following antibodies were used at the indicated dilutions: ATM (rabbit, Abcam, ab32420, 1/250), ATR (rabbit, Cell Signaling E1S3S 1/200), γH2AX (mouse, Millpore, 05-636, 1/500), CycA (mouse, BD biosciences, 611269, 1/100), MDC1 (mouse, Abcam, Ab50003, 1/300), 53BP1 (mouse, gift from T. Halazonetis, 1/20), TOPBP1 (A300-111A, Bethyl, 1/500 or rabbit, millipore, ABE1463, 1/250), NBS1 (rabbit, Novus, NB100-143, 1/200 or rabbit, Santa Cruz, sc-515069, 1/500), Treacle (rabbit, Sigma-Aldrich, HPA038237, 1/500).

### Widefield and confocal microscopy

Confocal images were acquired with a Leica SP8 inverse confocal laser scanning microscope coupled to a Leica DMI6000B inverted stand, with a 63X, 1.4-NA Plan-Apochromat oil-immersion objective. For triple-wavelength emission detection we combined DAPI with Alexa Fluor 488 or EGFP and Alexa Fluor 568. Images were taken using the sequential scanning mode and fields were recorded at a resolution of 512X512 pixels, 8-bit depth or 1024X1024 pixels, 8-bit depth. 3-10 z-sections were recorded with optimal distances based on Nyquist criterion. The maximum intensity projections were calculated, and images were adjusted for brightness and contrast in the figures. The Widefield image acquisition was done using three microscopes: first, a Zeiss AxioObserver.Z1 inverted widefield microscope, equipped with a Lumencor SpectraX illumination system and a Hamamatsu Orca Flash 4.0 V2, sCMOS cooled fluorescence camera (16bit, 2048 x 2048 pixel (4 MP), pixel size 6.5 µm). A 63x, 1.4-NA, i-plan apochromat oil-immersion objective was used. For triple-wavelength emission detection, we combined DAPI with Alexa fluor 488 or EGFP and Alexa Fluor 568. Second, a Leica DMI6000B inverted widefield microscope, equipped with Leica K5 sCMOS fluorescence camera (16-bit, 2048 x 2048-pixel, 4.2 MP) and Las X software version 3.7.2.22383. A HC Plan Apochromat 40X/0.95 PH dry objective and HCX Plan Apochromat 63X/1.40, and Plan Apochromat 100X/1.40 PH oil immersion objectives were used for image acquisition. Third, a Zeiss Axio Imager.Z2 equipped with a LED XCite 120LED illumination system and an ORCA-Flash4 LT Hamamatsu monochrome camera (16bit, 2048 x 2048 pixels, pixel size 6.5 µm). A 63X, 1.4 NA Plan Apochromat oil-immersion objective was used, with the ApoTome.2 engaged. For triple-wavelength emission detection, we combined DAPI with EGFP or Alexa Fluor 488 and Alexa Fluor 568. For optimal representation in figures, images were adjusted for brightness and exported as RGB TIF files using Fiji (Schindelin et al., 2012).

### Image quantification

The xyz confocal datasets (z-stacks) were analyzed using CellProfiler 4.2.8 (McQuin et al., 2018). First, nuclei segmentation was performed either by the intensity-based primary object detection module using the DAPI signal, or with the pretrained Cellpose “nuclei” model, within the “RunCellpose” CellProfiler plugin (Cellpose v3.0.9) (Stringer et al., 2021). For nucleolar caps and foci segmentation the primary object detection module was used on the respective channels after applying a feature enhancement filter. Downstream data manipulation and graphical representation of the data were done using Prism (GraphPad software) or R 3.4.2 (R Development Core Team). CellProfiler pipelines and R scripts are available upon request. For quantitative assessment of protein colocalization, the SQUASSH plugin (part of the MosaicSuite) for ImageJ and Fiji was used (Rizk et al., 2014). Within CellProfiler, foci detected in one fluorescence channel were masked and filtered against foci detected in the other channel, using the MaskObjects and FilterObjects modules to identify colocalized foci. The RelateObjects module was then used to assign these foci to their parent nuclei.

### Affinity capture of biotinylated proteins

Flp-In 293 T-REx cell lines stably expressing the recombinant opto-TOPBP1 protein were cultured to approximately 75% confluence and induced with doxycycline (2µg/mL) for 16 h. The following day, cells were incubated with biotin (500 µM) for 15 min in presence or absence of Blue Light as described (Frattini et al., 2021). Cells were subsequently washed with PBS and lysed in lysis buffer containing 50 mM Tris-HCl (pH 7.5), 150 mM NaCl, 1 mM EDTA, 1 mM EGTA, 1% NP-40, 0.2% SDS, and 0.5% sodium deoxycholate, supplemented with 1× complete protease inhibitor, 1× phosphatase inhibitor, and 250 U benzonase. Cell lysates were incubated on a rotating wheel for 1 h at 4°C and subsequently sonicated on ice (30% amplitude; three cycles of 10 s sonication followed by 2 s rest). Lysates were centrifuged at 7,750 × g for 30 min at 4°C, and the resulting clarified supernatants were transferred to fresh tubes. Total protein concentration was determined using the Bradford assay. For each condition, 1.5 mg of total protein was incubated with 50 µL of streptavidin-agarose beads on a rotating wheel for 3 h at 4°C. Following a 1-min centrifugation at 400 × g, the beads were sequentially washed with 1 mL of lysis buffer, 1 mL of Wash Buffer 1 (2% SDS in H₂O), 1 mL of Wash Buffer 2 (0.2% sodium deoxycholate, 1% Triton X-100, 500 mM NaCl, 1 mM EDTA, and 50 mM HEPES, pH 7.5), 1 mL of Wash Buffer 3 (250 mM LiCl, 0.5% NP-40, 0.5% sodium deoxycholate, 1 mM EDTA, 500 mM NaCl, and 10 mM Tris, pH 8), and 1 mL of Wash Buffer 4 (50 mM Tris, pH 7.5, and 50 mM NaCl). Bound proteins were eluted from the agarose beads using 40μl of 2X Laemmli Sample buffer and sent for mass spectrometry analysis. For Western blot analysis of TOPBP1 partners enriched in optogenetic TOPBP1 condensates, cells were simultaneously incubated with 500μM of biotin and exposed to blue light for 10 min of light-dark cycles (4 sec light followed by 30 sec dark). Biotin proximity labeling of light induced TOPBP1 partners was performed using streptavidin-coated beads as described previously. Bound proteins were eluted from the agarose beads with 80μl of 2X Laemmli sample buffer and incubated at 95°C for 10 min. 5μg of the lysates were used for Western blot analysis and probed by immunoblotting to detect proteins that are associated with TOPBP1 clusters, in the absence of DNA damage.

### Mass spectrometry

Sample digestion was essentially performed as described (Shevchenko et al., 2006). Briefly, proteins were loaded on a SDS-PAGE (BioRad, 456-1034) and, after short migration, a single band was excised. Proteins in the excised band were digested with Trypsin (Promega, V5280). The resulting peptides were analyzed online by nano-flow HPLC-nanoelectrospray ionization using a Qexactive HFX mass spectrometer (Thermo Fisher Scientific) coupled to a nano-LC system (Thermo Fisher Scientific, U3000-RSLC). Desalting and preconcentration of samples were performed online on a Pepmap! precolumn (0.3 3 10mm; Fisher Scientific, 164568). A gradient consisting of 0% to 40% B in A (A: 0.1% formic acid [Fisher Scientific, A117], 6% acetonitrile [Fisher Scientific, A955], in H2O [Fisher Scientific, W6], and B: 0.1% formic acid in 80% acetonitrile) for 120min at 300nl/min was used to elute peptides from the capillary reverse-phase column (0.075 3 250mm, Pepmap!, Fisher Scientific, 164941). Data were acquired using the Xcalibur software (version 4.0). A cycle of one full-scan mass spectrum (375–1,500 m/z) at a resolution of 60000 (at 200 m/z) followed by 12 data-dependent MS/MS spectra (at a resolution of 30000, isolation window 1.2 m/z) was repeated continuously throughout the nanoLC separation. Raw data analysis was performed using the MaxQuant software (version 1.5.5.1) with standard settings. The database used consisted of human entries from Uniprot (reference proteome UniProt 2018_09) and 250 contaminants (MaxQuant contaminant database).

### Flow cytometry (cell cycle analysis)

Cell cycle assays using BrdU and propidium iodide were performed as previously described in (Egger et al., 2022). Briefly, cells were pulse-labeled with 30µM BrdU for 15 min, harvested, washed in 1 × PBS and fixed by the dropwise addition of ice-cold ethanol to a final concentration of 70% (v/v). Cells were digested in a 30 mM HCl solution containing 0.5 mg/ml of pepsin, after which DNA was denatured with 2N HCl. Incorporated BrdU was immunodetected using a rat monoclonal anti-BrdU antibody (clone BU1/75 (ICR1)), followed by an Alexa Fluor 488-conjugated secondary antibody. DNA was stained with 25 µg/ml propidium iodide (PI) in 1 × PBS. Samples were acquired on a Gallios Flow Cytometer (Beckman Coulter). Debris and doublets were excluded based on forward- and side-scatter parameters (FSC/SSC) and PI height versus area. A total of 20,000 gated singlets were analyzed per biological condition, using the Kaluza dedicated software (Beckman Coulter).

---

**Supplementary Figure S1. Related to Figure 1.**
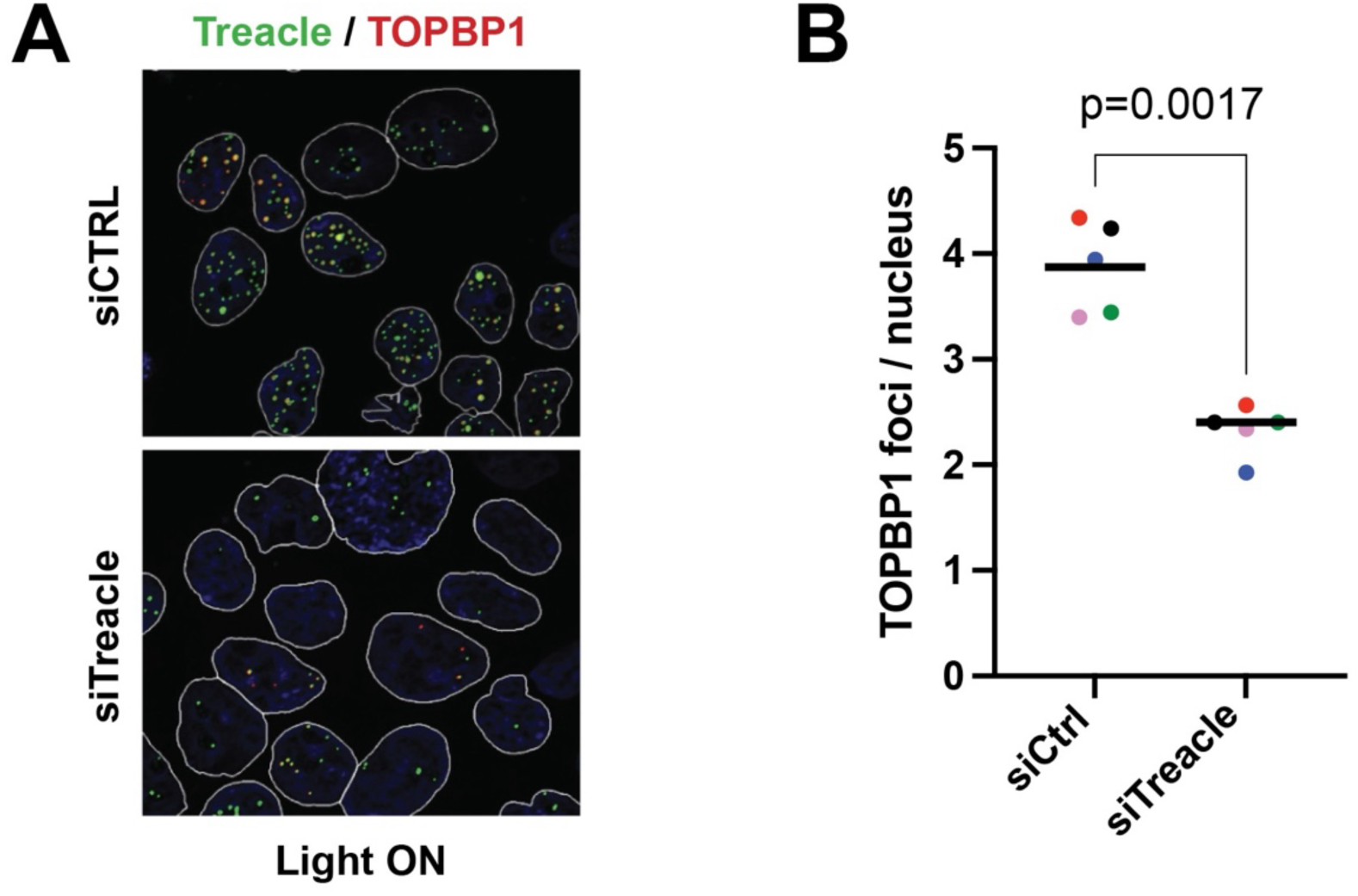
(A) Immunofluorescence staining with the indicated antibodies of opto-TOPBP1 expressing cells activated by light in siControl and siTreacle transfected cells. (B) Quantification of the experiment in (A). Dots show the mean number of TOPBP1 foci per nucleus in 5 independent experiments (color coded). Bar shows the mean of means. Statistical significance was tested by paired two-tailed t-test. Actual p-value is indicated (n=5; α=0.05).

**Supplementary Figure S2. Related to Figure 2.**
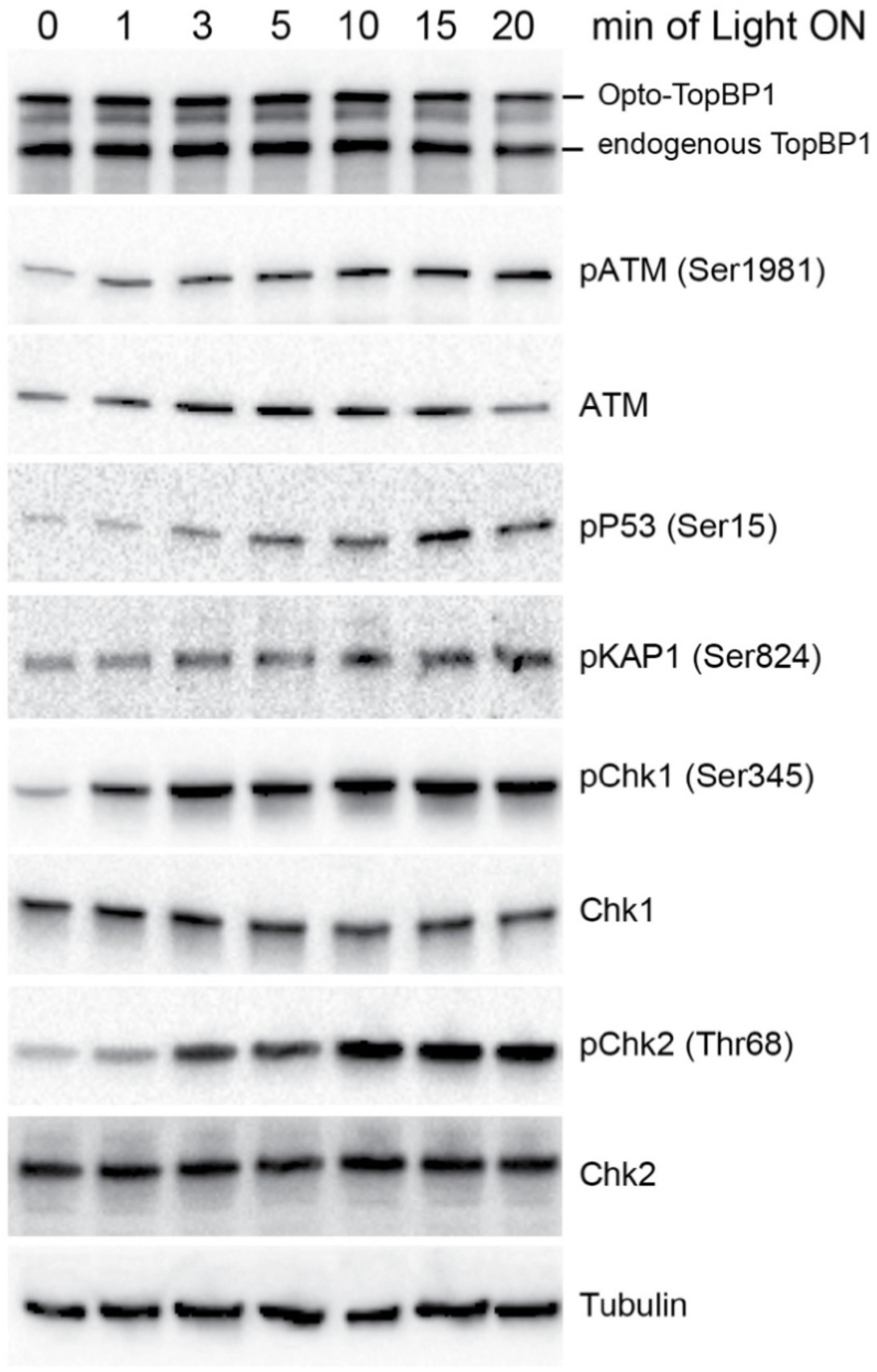
The indicated proteins were probed using immunoblotting cells expressing opto-TOPBP1 after light activation at various time points, as indicated.

**Supplementary Figure S3. Related to Figure 3.**
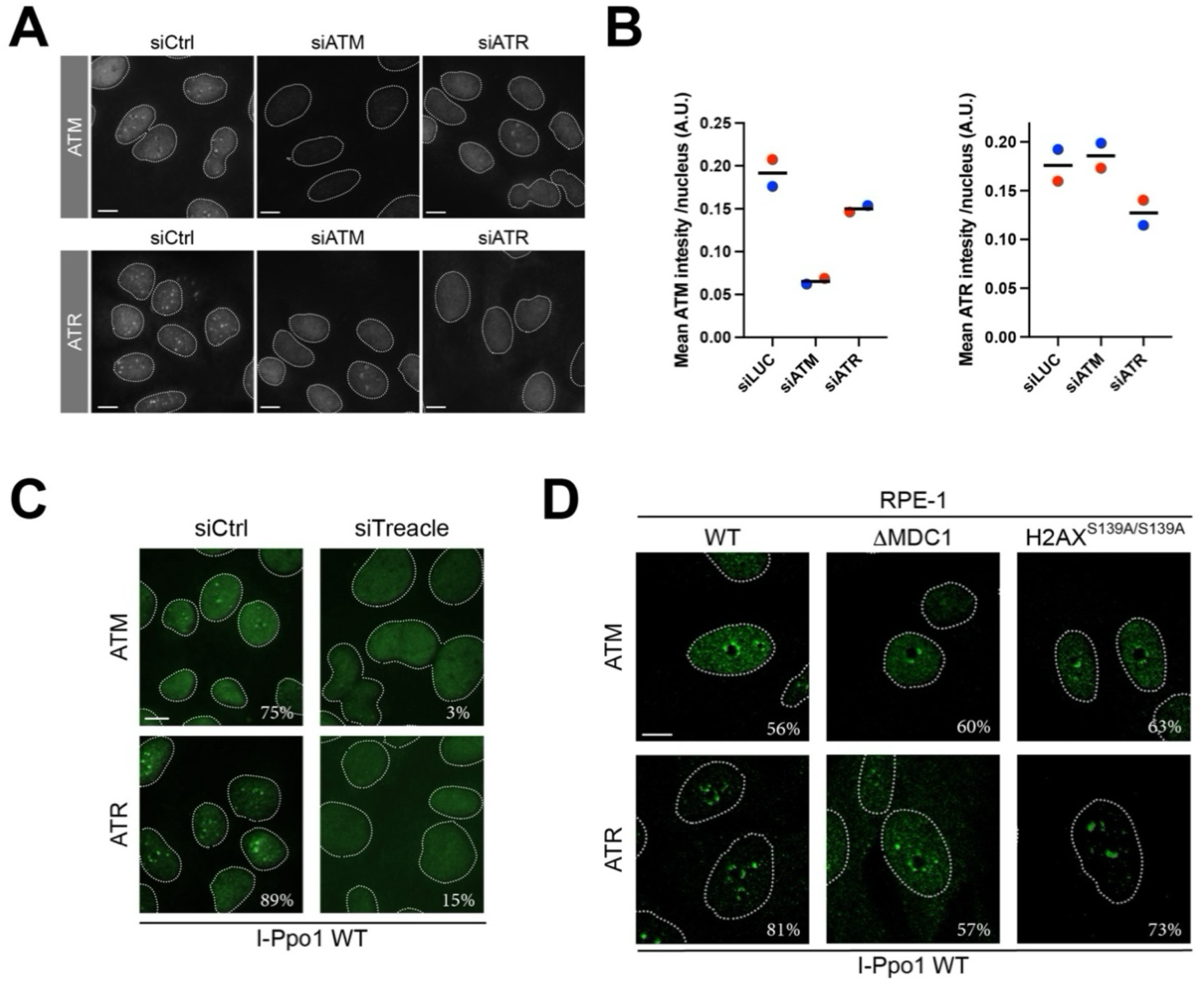
(A) ATM and ATR localization and staining intensity in ATM- and ATR-depleted U2OS cells and control cells 2h after I-PpoI transfection. (B) Quantification of the experiment in (A). Mean ATM and ATR intensity per nucleus was measured. Dots show the mean of two independent experiments (color coded) and black lines the mean of means. (C) ATM and ATR localization in Treacle-depleted U2OS cells and control cells 2h after I-PpoI transfection. Percentage of cells with > 2 ATM or ATR caps are indicated. (D) ATM and ATR localization in RPE1 ΔMDC1 cells, RPE1 H2AX^S139A/S139A^ cells and control cells 2h after I-PpoI transfection. Percentage of cells with > 2 ATM or ATR caps are indicated.

**Supplementary Figure S4. Related to Figure 5.**
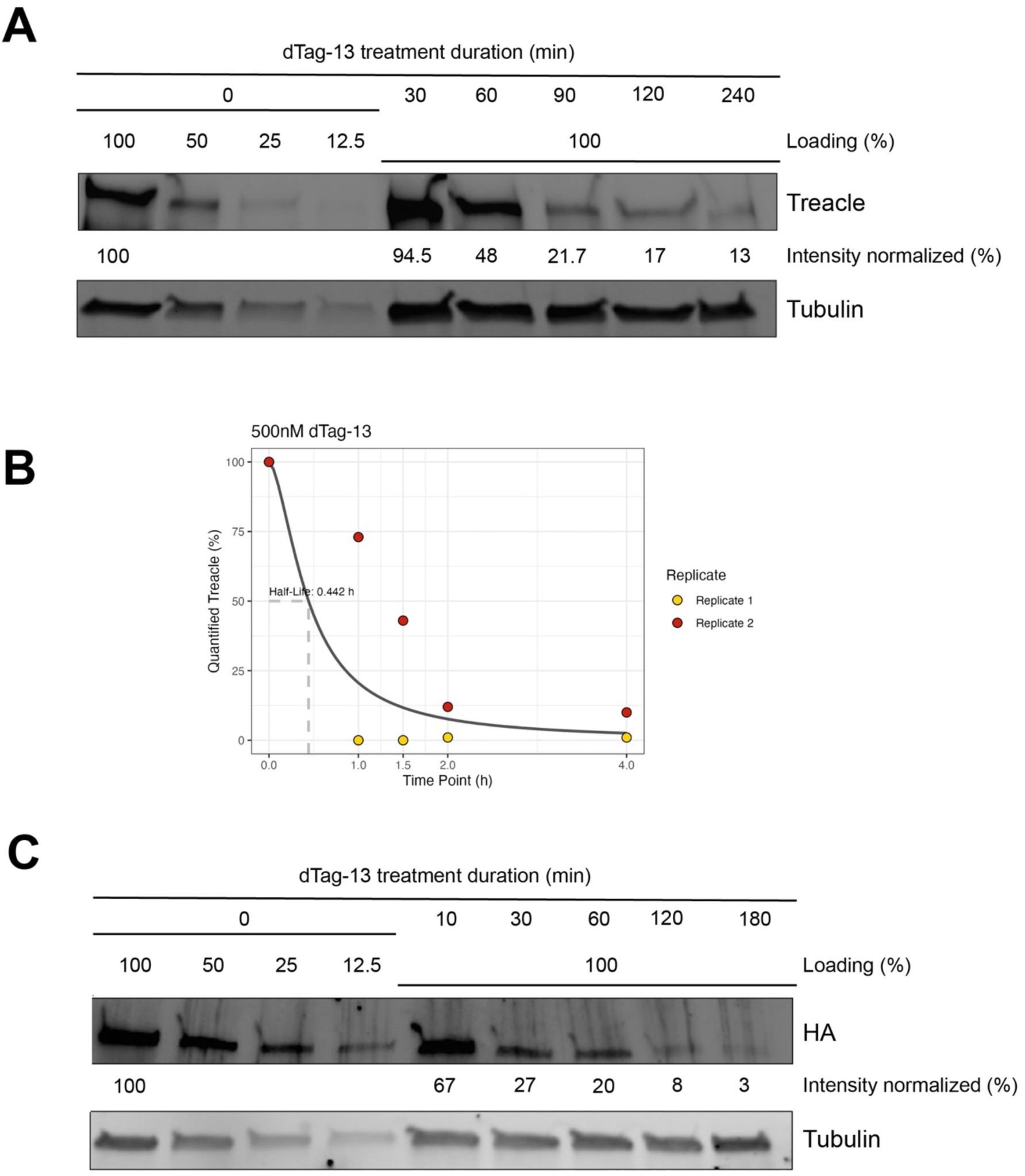
(A) Treacle^dTag^ depletion efficiency probed by Western blotting. Timepoints after dTag-13 addition are indicated. Relative protein levels are indicated. (B) Treacle^dTag^ half-life quantification upon dTag-13 treatment of two independent experiments. Treacle^dTag^ degradation kinetics were fitted by nonlinear least-squares regression using a four-parameter logistic model (L.4) implemented in the drc package in RStudio. (C) TOPBP1^dTag^ depletion efficiency probed by Western blotting. Timepoints after dTag-13 addition are indicated. Relative protein levels are indicated.

**Supplementary Figure S5. Related to Figure 6.**
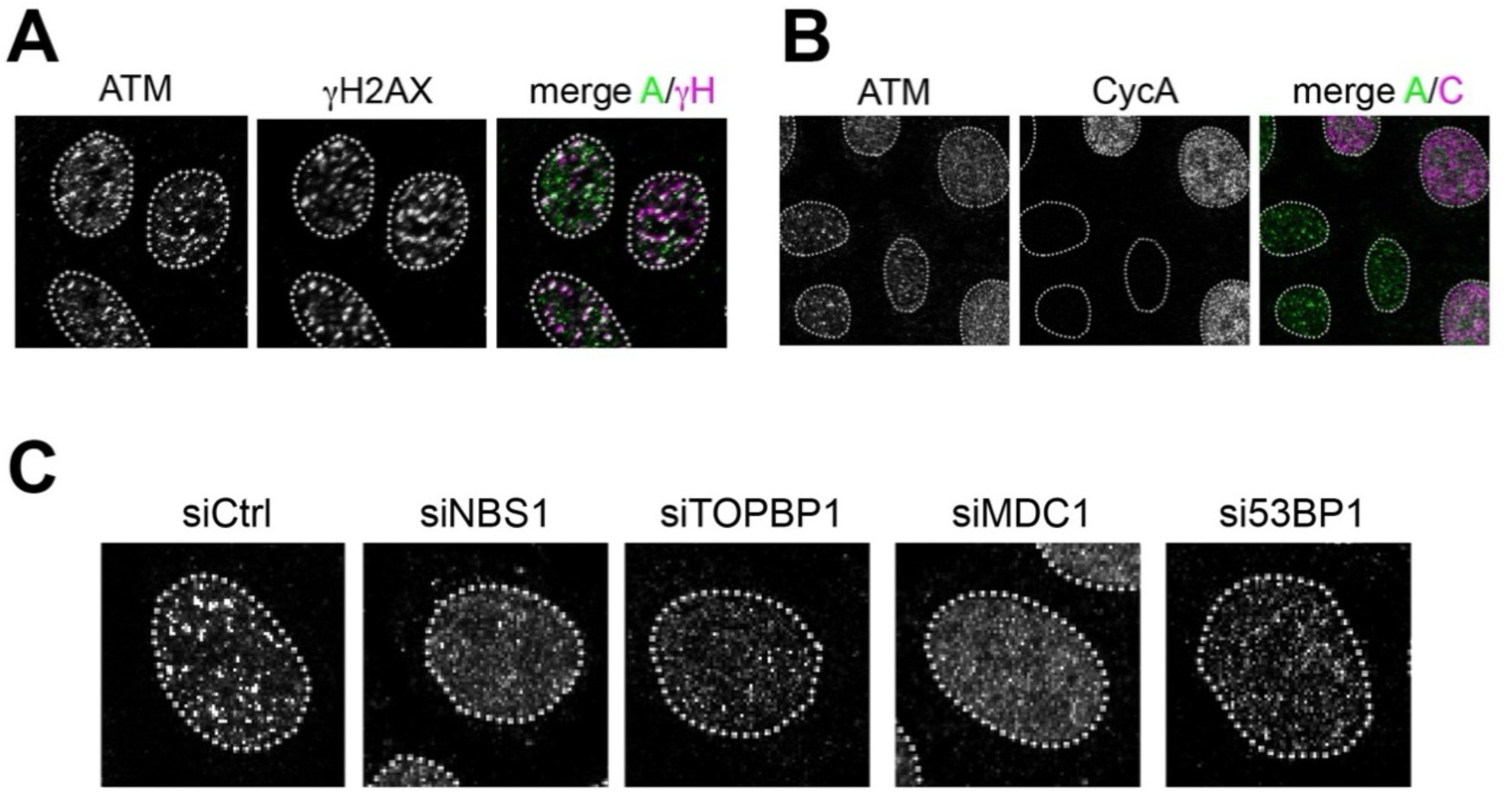
(A) Confocal micrographs of ATM and γH2AX localization in Hela cells 3h after exposure to 3Gy of IR. (B) Confocal micrographs of ATM and CyclinA (CycA) localization in Hela cells 3h after exposure to 3 Gy of IR. (C) Confocal micrographs of ATM localization in Hela cells either mock depleted (siCtrl) or depleted of NBS1 (siNBS1), TOPBP1 (siTOPBP1), MDC1 (siMDC1) and 53BP1 (si53BP1), respectively, 3h after exposure to 3 Gy of IR.

## References

Adam, S., Rossi, S.E., Moatti, N., De Marco Zompit, M., Xue, Y., Ng, T.F., Álvarez-Quilón, A., Desjardins, J., Bhaskaran, V., Martino, G. et al. (2021) The CIP2A-TOPBP1 axis safeguards chromosome stability and is a synthetic lethal target for BRCA-mutated cancer. Nat Cancer, 2, 1357–1371.

Blackford, A.N., and Jackson, S.P. (2017) ATM, ATR, and DNA-PK: The Trinity at the Heart of the DNA Damage Response. Molecular Cell, 66, 801–817.

Cescutti, R., Negrini, S., Kohzaki, M., and Halazonetis, T.D. (2010) TopBP1 functions with 53BP1 in the G1 DNA damage checkpoint. The EMBO Journal, 29, 3723–3732.

Chapman, J.R., and Jackson, S.P. (2008) Phospho-dependent interactions between NBS1 and MDC1 mediate chromatin retention of the MRN complex at sites of DNA damage. EMBO reports, 9, 795–801.

Chiang, T.W., le Sage, C., Larrieu, D., Demir, M., and Jackson, S.P. (2016) CRISPR-Cas9(D10A) nickase-based genotypic and phenotypic screening to enhance genome editing. Sci Rep, 6, 24356.

De Marco Zompit, M., Esteban, M.T., Mooser, C., Adam, S., Rossi, S.E., Jeanrenaud, A., Leimbacher, P.A., Fink, D., Shorrocks, A.K., Blackford, A.N., et al. (2022) The CIP2A-TOPBP1 complex safeguards chromosomal stability during mitosis. Nat Commun, 13, 4143.

Doil, C., Mailand, N., Bekker-Jensen, S., Menard, P., Larsen, D.H., Pepperkok, R., Ellenberg, J., Panier, S., Durocher, D., Bartek, J. et al. (2009) RNF168 binds and amplifies ubiquitin conjugates on damaged chromosomes to allow accumulation of repair proteins. Cell, 136, 435–446.

Egger, T., Morano, L., Blanchard, M.P., Basbous, J., and Constantinou, A. (2024) Spatial organization and functions of Chk1 activation by TopBP1 biomolecular condensates. Cell Rep, 43, 114064.

Falck, J., Coates, J., and Jackson, S.P. (2005) Conserved modes of recruitment of ATM, ATR and DNA-PKcs to sites of DNA damage. Nature, 434, 605–611.

Frattini, C., Promonet, A., Alghoul, E., Vidal-Eychenie, S., Lamarque, M., Blanchard, M.P., Urbach, S., Basbous, J., and Constantinou, A. (2021) TopBP1 assembles nuclear condensates to switch on ATR signaling. Mol Cell, 81, 1231–1245.e8.

Huen, M., Grant, R., Manke, I., Minn, K., Yu, X., Yaffe, M., and Chen, J. (2007) RNF8 Transduces the DNA-Damage Signal via Histone Ubiquitylation and Checkpoint Protein Assembly. Cell, 131, 901–914.

Kolas, N.K., Chapman, J.R., Nakada, S., Ylanko, J., Chahwan, R., Sweeney, F.D., Panier, S., Mendez, M., Wildenhain, J., Thomson, T.M. et al. (2007) Orchestration of the DNA-Damage Response by the RNF8 Ubiquitin Ligase. Science (New York, NY), 318, 1637–1640.

Larsen, D.H., Hari, F., Clapperton, J.A., Gwerder, M., Gutsche, K., Altmeyer, M., Jungmichel, S., Toledo, L.I., Fink, D., Rask, M.-B. et al. (2014) The NBS1-Treacle complex controls ribosomal RNA transcription in response to DNA damage. Nature cell biology, 16, 792–803.

Leimbacher, P.-A., Jones, S.E., Shorrocks, A.-M.K., de Marco Zompit, M., Day, M., Blaauwendraad, J., Bundschuh, D., Bonham, S., Fischer, R., Fink, D. et al. (2019) MDC1 Interacts with TOPBP1 to Maintain Chromosomal Stability during Mitosis. Molecular Cell, 74, 571–583.e8.

Lim, Y., Tamayo-Orrego, L., Schmid, E., Tarnauskaite, Z., Kochenova, O.V., Gruar, R., Muramatsu, S., Lynch, L., Schlie, A.V., Carroll, P.L. et al. (2023) In silico protein interaction screening uncovers DONSON’s role in replication initiation. Science, 381, eadi3448.

Lindsey-Boltz, L.A., and Sancar, A. (2011) Tethering DNA Damage Checkpoint Mediator Proteins Topoisomerase II -binding Protein 1 (TopBP1) and Claspin to DNA Activates Ataxia-Telangiectasia Mutated and RAD3-related (ATR) Phosphorylation of Checkpoint Kinase 1 (Chk1). Journal of Biological Chemistry, 286, 19229–19236.

Mailand, N., Bekker-Jensen, S., Faustrup, H., Melander, F., Bartek, J., Lukas, C., and Lukas, J. (2007) RNF8 ubiquitylates histones at DNA double-strand breaks and promotes assembly of repair proteins. Cell, 131, 887–900.

McQuin, C., Goodman, A., Chernyshev, V., Kamentsky, L., Cimini, B.A., Karhohs, K.W., Doan, M., Ding, L., Rafelski, S.M., Thirstrup, D. et al. (2018) CellProfiler 3.0: Next-generation image processing for biology. PLoS Biology, 16, e2005970–17.

Melander, F., Bekker-Jensen, S., Falck, J., Bartek, J., Mailand, N., and Lukas, J. (2008) Phosphorylation of SDT repeats in the MDC1 N terminus triggers retention of NBS1 at the DNA damage-modified chromatin. J Cell Biol, 181, 213–226.

Mooser, C., Symeonidou, I.E., Leimbacher, P.A., Ribeiro, A., Shorrocks, A.K., Jungmichel, S., Larsen, S.C., Knechtle, K., Jasrotia, A., Zurbriggen, D. et al. (2020) Treacle controls the nucleolar response to rDNA breaks via TOPBP1 recruitment and ATR activation. Nat Commun, 11, 123.

Nabet, B., Roberts, J.M., Buckley, D.L., Paulk, J., Dastjerdi, S., Yang, A., Leggett, A.L., Erb, M.A., Lawlor, M.A., Souza, A. et al. (2018) The dTAG system for immediate and target-specific protein degradation. Nat Chem Biol, 14, 431–441.

Park, A., Won, S.T., Pentecost, M., Bartkowski, W., and Lee, B. (2014) CRISPR/Cas9 Allows Efficient and Complete Knock-In of a Destabilization Domain-Tagged Essential Protein in a Human Cell Line, Allowing Rapid Knockdown of Protein Function. PloS one, 9, e95101.

Petrova, N.V., Deriglazov, D.A., Kovina, A.P., Luzhin, A.V., Shender, V.O., Arapidi, G.P., Gavrikov, A.S., Mishin, A.S., Razin, S.V., and Velichko, A.K. (2026) Treacle-dependent TOPBP1 condensation regulates the nucleolar DNA damage response. Nucleic Acids Res, 54, gkag350.

Rizk, A.E.L., Paul, G.E.G., Pietro, I., Bugarski, M., Mansouri, M., Niemann, A., Ziegler, U., Berger, P., and Sbalzarini, I.F. (2014) Segmentation and quantification of subcellular structures in fluorescence microscopy images using Squassh. Nature Protocols, 9, 586–596.

Schindelin, J., Arganda-Carreras, I., Frise, E., Kaynig, V., Longair, M., Pietzsch, T., Preibisch, S., Rueden, C., Saalfeld, S., Schmid, B. et al. (2012) Fiji: an open-source platform for biological-image analysis. Nature Methods, 9, 676–682.

Shevchenko, A., Tomas, H., Havlis, J., Olsen, J.V., and Mann, M. (2006) In-gel digestion for mass spectrometric characterization of proteins and proteomes. Nat Protoc, 1, 2856–2860.

Sokka, M., Rilla, K., Miinalainen, I., Pospiech, H., and Syvaoja, J.E. (2015) High levels of TopBP1 induce ATR-dependent shut-down of rRNA transcription and nucleolar segregation. Nucleic acids research, 43, 4975–4989.

Spycher, C., Miller, E.S., Townsend, K., Pavic, L., Morrice, N.A., Janscak, P., Stewart, G.S., and Stucki, M. (2008) Constitutive phosphorylation of MDC1 physically links the MRE11-RAD50-NBS1 complex to damaged chromatin. J Cell Biol, 181, 227–240.

Stewart, G.S., Panier, S., Townsend, K., Al-Hakim, A.K., Kolas, N.K., Miller, E.S., Nakada, S., Ylanko, J., Olivarius, S., and Mendez, M. (2009) The RIDDLE Syndrome Protein Mediates a Ubiquitin-Dependent Signaling Cascade at Sites of DNA Damage. Cell, 136, 420–434.

Stringer, C., Wang, T., Michaelos, M. and Pachitariu, M. (2021) Cellpose: a generalist algorithm for cellular segmentation. Nat Methods, 18, 100–106.

Trivedi, P., Steele, C.D., Au, F.K.C., Alexandrov, L.B., and Cleveland, D.W. (2023) Mitotic tethering enables inheritance of shattered micronuclear chromosomes. Nature, 618, 1049–1056.

van Sluis, M., and McStay, B. (2015) A localized nucleolar DNA damage response facilitates recruitment of the homology-directed repair machinery independent of cell cycle stage. Genes Dev, 29, 1151–1163.

Velichko, A.K., Petrova, N.V., Luzhin, A.V., Strelkova, O.S., Ovsyannikova, N., Kireev, I.I., Petrova, N.V., Razin, S.V., and Kantidze, O.L. (2019) Hypoosmotic stress induces R loop formation in nucleoli and ATR/ATM-dependent silencing of nucleolar transcription. Nucleic Acids Research, 47, 6811–6825.

Wang, Y.L., Zhao, W.W., Bai, S.M., Feng, L.L., Bie, S.Y., Gong, L., Wang, F., Wei, M.B., Feng, W.X., Pang, X.L. et al. (2022) MRNIP condensates promote DNA double-strand break sensing and end resection. Nat Commun, 13, 2638.

Wardlaw, C.P., Carr, A.M., and Oliver, A.W. (2014) TopBP1: A BRCT-scaffold protein functioning in multiple cellular pathways. DNA Repair, 22, 165–174.

Warren, C., and Pavletich, N.P. (2022) Structure of the human ATM kinase and mechanism of Nbs1 binding. Elife, 11, e74218.

Wu, L., Luo, K., Lou, Z., and Chen, J. (2008) MDC1 regulates intra-S-phase checkpoint by targeting NBS1 to DNA double-strand breaks. Proceedings of the National Academy of Sciences of the United States of America, 105, 11200–11205.

